# “Estimating abundance and phenology from transect count data with GLMs”

**DOI:** 10.1101/2020.06.01.127910

**Authors:** Collin Edwards, Elizabeth E. Crone

**Affiliations:** Department of Biology, Tufts University, Medford MA 02155 USA

**Keywords:** population dynamics, phenology, climate change, peak abundance, activity period, first emergence, Gaussian curve, Euphydryas phaeton, growing degree days, general linear model

## Abstract

Estimating population abundance is central to population ecology. With increasing concern over declining insect populations, estimating trends in abundance has become even more urgent. At the same time, there is an emerging in interest in quantifying phenological patterns, in part because phenological shifts are one of the most conspicuous signs of climate change. Existing techniques to fit activity curves (and thus both abundance and phenology) to repeated transect counts of insects (a common form of data for these taxa) frequently fail for sparse data, and often require advanced knowledge of statistical computing. These limitations prevent us from understanding both population trends and phenological shifts, especially in the at-risk species for which this understanding is most vital. Here we present a method to fit repeated transect count data with Gaussian curves using linear models, and show how robust abundance and phenological metrics can be obtained using standard regression tools. We then apply this method to eight years of Baltimore checkerspot data using generalized linear models (GLMs). This case study illustrates the ability of our method to fit even years with only a few non-zero survey counts, and identifies a significant negative relationship between population size and annual variation in thermal environment (in growing degree days). We believe our new method provides a key tool to unlock previously-unusable sparse data sets, and may provide a useful middle ground between ad hoc metrics of abundance and phenology and custom-coded mechanistic models.

## Introduction

Ecologists are observing massively elevated extinction rates (Turvey and Crees 2019), driven in part by direct anthropogenic activities, climate change, and the spread of invasive species (Pievani 2014). We are also seeing frequent changes in the phenology of populations, a “globally coherent fingerprint of climate change” (Parmesan and Yohe 2003). Both of these patterns are particularly pronounced in insects, for which there are alarming signs of declining populations for many well-studied taxa (e.g. Thomas et al. 2005, Forister et al. 2010, Potts et al. 2010) and more broadly (e.g. Hallman et al. 2017, van Klink et al. 2020). However, evidence for global trends is mixed, with other studies showing no overall trends and in some cases contradicting previous papers that found declines (Wagner et al. 2021). This conflicting literature highlights the limitations of current tools and data sets (Thomas et al. 2019, Didham et al. 2020). One key limitation is often the lack of data to estimate trends for individual species or populations as opposed to broad taxonomic groups or guilds (Wagner et al. 2021), which is especially problematic for rare or at-risk species.

One of the common forms of sampling for insect populations are systematic repeated surveys throughout an activity period, such as “Pollard” transect walks (Pollard 1977), bee bowls (e.g. Stemkovski et al. 2020), or trap nests (Forrest and Thomson 2011). Historically, the main goal of these surveys was simply to estimate yearly abundance (e.g. Zonneveld 1991, Pollard and Yates 1993, Schultz and Hammond 2003). More recently, there has been growing interest in also estimating phenology from this type of data, starting at least with Sparks and Yates (1997), but with considerable recent interest (e.g. Stewart et al. 2020, Fric et al. 2020). Estimating abundance and phenology from repeated count data seems like it should be easy, yet often remains a challenge. Initial approaches for estimating abundance involved averaging the counts of surveys across the activity period (Pollard et al. 1975, Pollard 1977, Thomas 1983, Pollard and Yates 1993), which has clear limitations (e.g. requires appropriate estimation of activity period, appropriate sampling within activity period, and if the same population spreads its activity across a longer period, the average count will shrink). Initial approaches for estimating phenology often looked at the first day individuals were observed (e.g. Sparks and Yates 1997), but this metric can covary with population abundance and sampling effort, so can confound phenological shifts with other changes (Van Strien et al. 2008, Miller-Rushing et al. 2008, Inouye et al. 2019).

To improve on these basic approaches, numerous studies have proposed realistic or highly flexible models (for a list of examples, see Table 1). However, with few exceptions, these methods were developed or proposed in the context of repeated measures of flowering plants, where there are often dozens of time points in a year (e.g. Malo 2002, Clark and Thompson 2011, Malo 2002, Clark and Thompson 2011, Proia et al. 2015, Austen et al. 2014). Perhaps as a consequence of being developed with such rich data, current methods generally require considerable data to work. This limitation holds both for the suite of models developed for flowering plants, the “Zonneveld model” – a mechanistic phenology curve commonly used to analyze insect counts (Zonneveld 1991, INCA 2002, Haddad et al. 2008) --, and more generic approaches like generalized additive models (GAMs) (Rothery and Roy 2001, Hodgson et al. 2011, Newson et al. 2016, Stemkovski et al. 2020).

**Table 1:**
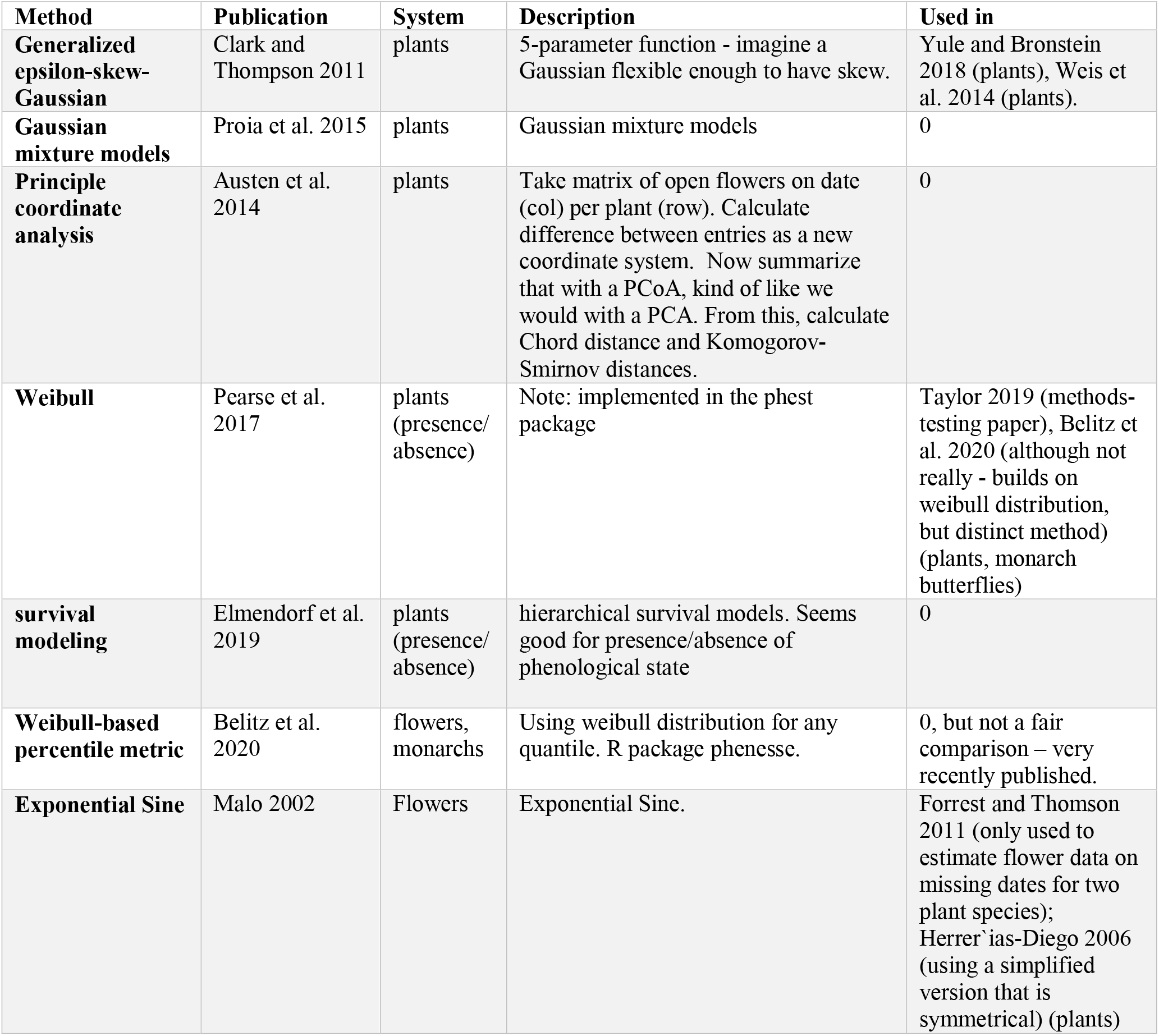
Summary of ad-hoc literature review on statistical methods for fitting activity curves to repeated count data, looking to see how often the proposed methods have actually been used. Note that there were a few non-English publications citing these methods papers which we were unable to evaluate. In addition to these methods, Generalized Additive Models (GAMs) have been widely used in a variety of phenological studies and one older method (Zonneveld et al. 1991) is widely used by some insect ecologists (see discussion in main text). We also did not include custom-coded (typically Bayesian) approaches that would need substantial recoding of the method to be applied to a new data set.

Ecologists sometimes address the limitations of current analytical techniques by working only with abundant species, or years for which there are many non-zero survey counts. For example, in a recent analysis of Ohio butterfly populations using GAMs, Wepprich et al. (2019) limited their analysis to cases where they had 10 or more surveys in a year. When fitting Spanish butterfly populations with Gaussian curves using a Bayesian model, Stewart et al. (2020) use only species that were present in at least half of their surveys, with at least 35 individuals observed per year. In their analysis of UK butterfly populations using GAMs, Hodgson et al. (2011) generally excluded sites where the species was observed in less than half the surveys. In simulations of data for the rare St. Francis’ satyr butterfly, Haddad et al. (2008) found that when survey frequency dropped to three times per week, the Zonneveld model (implemented using INCA (2002)) failed more than 30% of the time.

While ecologists can gain a lot of information by fitting elegant models to rich data sets, having only tools that require rich data may prevent the analysis of rare species or years of low abundance, both of which are likely to lead to infrequent non-zero survey counts. Ignoring rare species in turn can bias our understanding of global trends (Didham et al. 2020), and ignoring years of low abundance limits our ability to infer population dynamics or carry out population viability analysis (e.g. Gerber and Demaster 1999, Morris et al. 2002). To make matters worse, even with considerable data, there is no guarantee that existing methods can be solved numerically. For example, the Zonneveld model, which has become something of a standard for Pollard-walk style time series, can run into issues of confounded parameters; it is difficult to tell if you have a few long-lived butterflies or many short-lived ones, leading the Zonneveld model to fail (Gross et al. 2007, Table S1). Similarly, Malo (2002) presented an elegant phenological model based on the exponential sine function, but found that their numerical solvers failed to find reasonable solutions. They thus had to modify the 5-parameter model to include two additional parameters per year – defining the beginning and ending of the activity peaks for each year – which have to be determined ad-hoc by users for each year of data. In some other cases, Bayesian methods are recommended when data are sparse compared to model complexity. However, custom-coded Bayesian analyses can fail in ways that are not obvious to non-experts (Lele and Dennis 2009, Seaman et al. 2012).

A final challenge with current analytical methods is that many require substantial knowledge of computational statistics to implement successfully. There are certainly statistical ecologists with the skill and experience to write custom-coded hierarchical Bayesian models and ensure that the resulting estimates are sensible (e.g. Lindén and Mäntyniemi 2011, Chapman et al. 2015), but they are the minority of ecologists. Of the statistical methods we encountered in writing this paper, only two (the Zonneveld model and GAMs) have seen much use. Not coincidentally, these are the two methods with easy-to-use program implementations (INCA (INCA 2002) and the **mgcv** package in R (Wood 2017), respectively). In contrast, another seven methods published in the last 20 years have only been used in subsequent publications a combined total of six times^1^ (Table 1), and only one of those was applied to insect data (Belitz et al. (2020), itself proposing a new method). To date, the majority of the apparent surplus of analytical tools for repeated count data are not actually being used to study insect abundance or phenology.

Taken together, it is clear that ecologists lack an accessible, robust statistical tool for quantifying population abundance and phenology for species, years, or sites with sparse data. In this paper, we propose an approach to fit such data with Guassian curves using generalized linear models (GLMs). To illustrate this method, we first outline the algebra behind the procedure, then demonstrate its application to a 9-year time series of monitoring from a population of Baltimore checkerspot butterflies (*Euphydryas phaeton*) in Massachusetts. In the supplements we offer a detailed explanation of how to implement this approach in the programming language R (R Core Team 2020), and provide simple code to act as a template. The simplicity of Gaussian curves (defined by only 3 parameters) means that our proposed method can be applied to almost all data – we find that even three non-zero days of count is sufficient to fit an activity curve (admittedly one with wide confidence intervals). The familiarity of linear regression and Gaussian curves (and our example code) make this approach accessible to any ecologist who can run a linear regression in R.

## Gaussian curve as a linear model

The basis of our method is that a Gaussian curve has the form

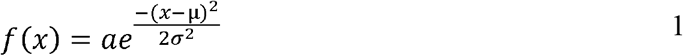

When *a* is chosen to make Equation 1 integrate to 1, this is the normal or Gaussian distribution. Since everything in Equation 1 is a constant except for *x*, if we multiply it out and define β_0_, β_1_, β_2_ appropriately in terms of the other constants (see Appendix S1 for the algebra), we can rewrite the Gaussian curve as

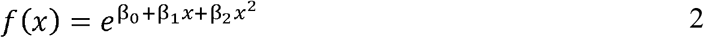

Here we can see that the terms in the exponent are a quadratic equation. This means that if we take the natural log of both sides, we are left with a familiar linear model with both a linear and a quadratic term:

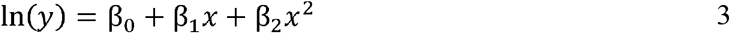

(Note that to produce an appropriate Guassian curve, β_1_ must be positive and β_2_ must be negative; otherwise this equation produces a monotonic or convex curve). Despite our special use for it, Equation 3 is an ordinary linear model of a quadratic equation, and can be fit with standard tools for linear models. In the context of phenology, the most straightforward analysis would use empirical estimates of abundance or activity (e.g. transect counts of butterflies or flowers) for dependent variable *y*, which is distributed following a Gaussian curve in relation to some measure of time (e.g. day of year) for the independent variable *x* (Fig. 1A-I).

**Figure 1:**
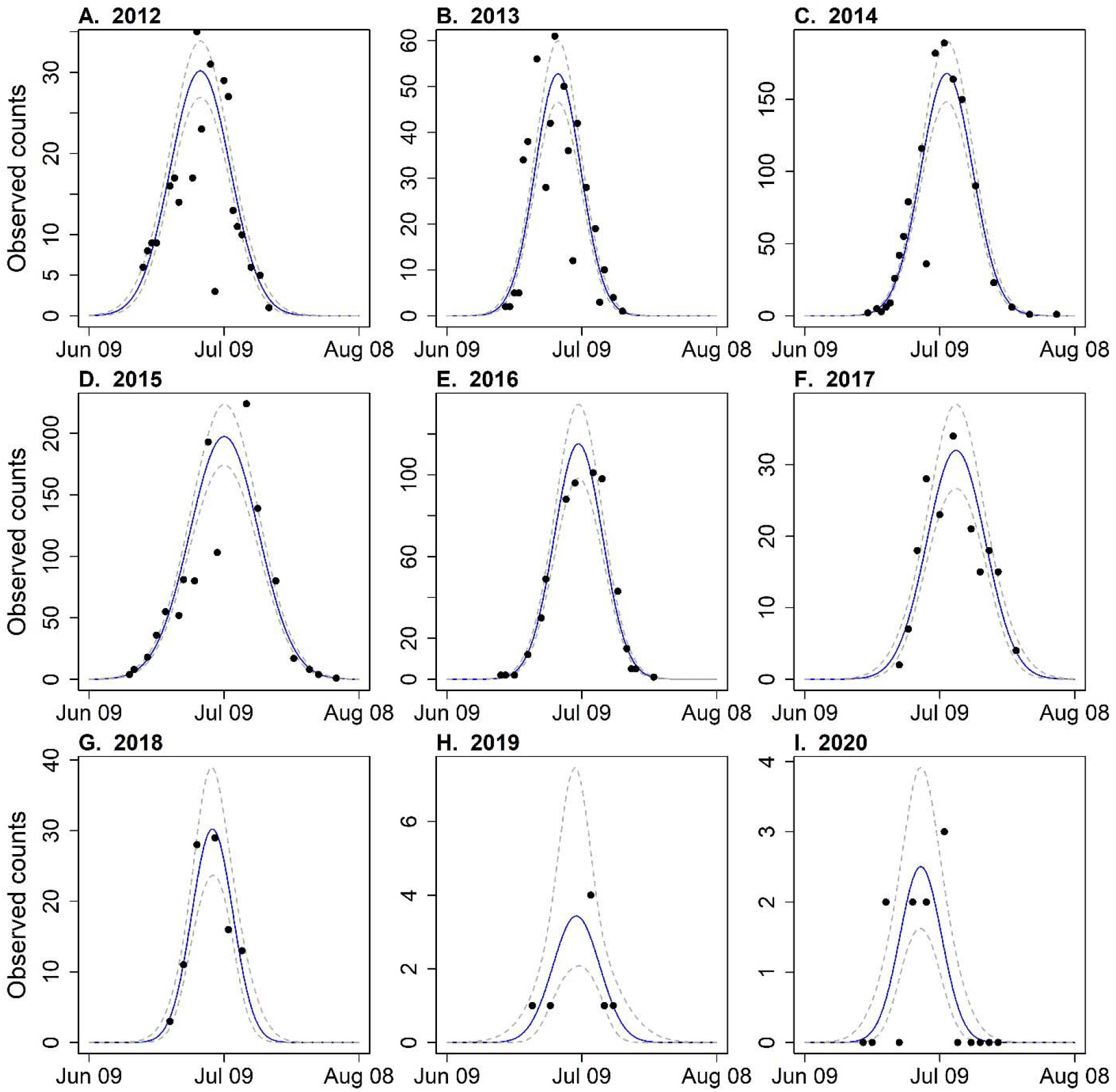
Gaussian model fitted to Baltimore checkerspot butterfly data. Points show raw data, blue lines show best-fitting Gaussian curve, dashed gray lines show +/− 1 standard error. For comparability, day of month for axis labels in this and other figures is based on a 365 day year (excludes leap days). Note the different scales on the y-axes.

Fitting a linear model of *ln(y)* vs. *x* provides estimates and confidence intervals for β_0_, β_1_, the Gaussian curve (mean *μ,* variance *σ*^2^), as well as metrics determined by the gaussian curve and β_2_. By reversing the algebra between Equations 1 and 2, we can recover the parameters of (e.g. area under the curve) that may be useful in interpreting the fitted activity curves (Appendix S1: Fig. S1). First, *μ,* the estimated **day of peak activity** and **mean day of activity** (these are the same since the Gaussian curve is symmetrical) can be calculated from the slopes of the linear and quadratic terms:

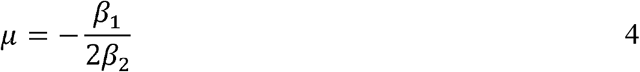

The standard deviation of the Gaussian curve, *σ*, is a function of the slope of the quadratic term:

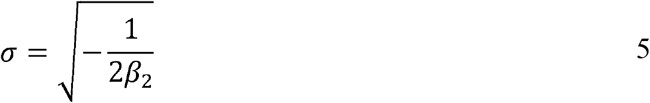

In the case study below, we refer to 2*1.285\**σ* as the **activity period**. This corresponds to the range of dates between the 0.1 and the 0.9 quantiles, and so this measures the duration of time when the middle 80% of observations are estimated to occur (e.g. Jonzén et al. 2006, Michielini et al. 2020).

The area under the gaussian curve, N, is a **population abundance index:**

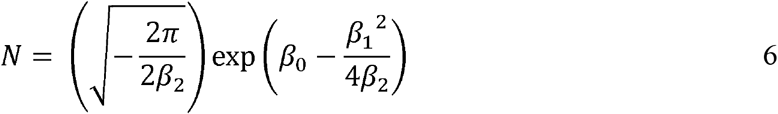

We use the term “abundance index” because, for constant sampling effort, survival, and detection probability, *N* will be proportional to the number of active individuals (e.g. Gross et al. 2007). However, it is actually a measure of estimated observed activity-days (Dennis et al. 2015, Wepprich et al. 2019).

We note that here we have focused on a few phenological metrics, including the days of 0.1, 0.5 (e.g. day of peak) and 0.9 quantile. However, because our proposed approach fits a Guassian curve, from estimated μ and *σ* it is trivial to calculate any characteristics of a Gaussian curve, including (a) any arbitrary quantiles (e.g. the 0.05 and 0.95 quantiles used in Stemkovski et al. 2020), (b) the height of the curve (e.g. maximum number of flowers, Miller-Rushing and Inouye (2009)), or (c) the “observable flight season” (days when the curve exceeds 1, a metric reflecting the period of likely human detection that parallels first and last observation dates) (Bonoan et al., *in review*).

Standard errors of derived parameters such as *μ*, *σ*^2^, or *N*, can be estimated using the delta method (Williams et al. 2002), or by parametric bootstrapping (Dennis 1996). Code for these analyses is given in Appendices S2 (tutorial as html), S3 (analysis as html) and S4 (data and Rmarkdown sources for Appendices S2 and S3). All code was written and run in R version 4.0.0 (R Core Team, 2020).

## Case study

### Data set

From 2012-2020, we conducted a capture-recapture study of Baltimore checkerspot (*Euphydryas phaeton*) butterflies at a natural area (Williams Conservation Land) in the town of Harvard MA, USA (Brown and Crone 2016, Brown et al. 2017, Crone 2018). Baltimore checkerspot is a univoltine species, with one clear flight period of adults per year. Surveys were conducted by visiting the site 2-3 times a week from mid-June until the population was clearly finished for the year; the onset of checkerspot flight at this site is usually in late June or early July. To illustrate the use of a Gaussian curve to estimate phenological metrics, we converted capture-recapture data to counts of individual animals handled on each visit to the site. This monitoring protocol creates a data structure that is similar to traditional “Pollard walk” style monitoring (Pollard 1977, Pollard and Yates 1993, Wepprich et al. 2019) but differs from Pollard walks in that the site was searched freely, rather than by walking a fixed route. For comparison with the Gaussian analyses below, we estimated population size each year using standard open population capture-recapture models (see supplemental methods). For comparison to existing methods, we fit our data to the Zonneveld model using INCA (INCA 2002). We chose to compare with the Zonneveld model because it is also easy to use, and as a 4-parameter model, it is one of the simplest (and thus most likely to fit our sparse data).

### Methods

#### Estimation of phenology metrics

We fit Gaussian curves to these data using generalized linear models (GLM) with a negative binomial family and log link function, with the number of butterflies seen on each day as the dependent variable, day of year and day of year squared as independent variables; we used a single model with interaction terms to allow curves to fit each year separately, but with shared estimation of the model variance term. After fitting the linear model, we used equations 4-7 to calculate the estimated mean day of activity, standard deviation of flight period, population abundance, and peak abundance for each year, and use mean day of activity and standard deviation of activity to calculate onset (day of 0.1 quantile) and end (day of 0.9 quantile) of activity. For heuristic purposes, we calculated confidence intervals for these metrics using both the delta method and parametric bootstrapping. To test for asymmetry in activity curves (a feature common in some systems and models), we regressed residuals by day of year using a linear model and then again with a cubic regression spline (using the **mgcv** package) (Wood 2017).

#### Comparison to INCA fits

We used the INCA program to fit each year of our data (INCA 2002). We carried out this analysis with INCA twice: first we fit INCA using default settings, putting INCA on a level playing field with the Gaussian method; second, we fit INCA again, providing an informative prior on mortality rate, defined by the mean and standard error of daily mortality for the Baltimore Checkerspot (Brown and Crone 2016, their Table 1). Both INCA and the Gaussian curve produce indices of population abundance rather than complete population estimates, and these indices are on different scales. As such, we focus on the correlation between INCA and Gaussian metrics.

#### Evaluating an environmental driver

After estimating abundance and phenology, a common next question is to ask whether changes in these population characteristics are associated with changes in environmental conditions (see, e.g., Roy and Sparks 2000, Forister and Shapiro 2003, Marra et al. 2005, Jonzén et al. 2006, Miller-Rushing et al. 2008, van Buskirk et al. 2009, Hodgson et al. 2011, Gordo et al. 2013, Bertin 2015, Cayton et al. 2015, Barton and Sandercock 2018, Heberling et al. 2019, Oke et al. 2019, Park et al. 2019, Fric et al. 2020, Horton et al. 2020, Stewart et al. 2020, Stemkovski et al. 2020). It is possible in principle to simultaneously fit drivers of population or phenological change and parameters themselves (e.g. Mizel et al. 2019). However, the algebra of converting a Gaussian curve to a linear model does not enable easy inclusion of covariates of the ecologically meaningful derived metrics such as onset of activity or peak dates (XXX, unpubl. calculations). One accessible alternative to custom-coding complex models is a two-step process of first estimating derived parameters (e.g. population abundance index) with linear models, then using these derived parameters in subsequent models (e.g. the approach used in Wepprich et al. (2019), but using GLMs instead of GAMs for the first step). To account for uncertainty in derived parameters, it is straightforward to use parameteric bootstrapping.

To illustrate this approach, we compared yearly estimates of the day of mean activity, flight period, and population abundance to temperature. Determining the most appropriate metrics to capture environmental drivers of population dynamics or phenology is an open question in ecology, and beyond the scope of this study. We instead chose to demonstrate the principles with growing degree days (GDD), a common measure of thermal environment that has been found to predict plant and insect phenology (e.g. Hodgson et al. 2011, Cayton et al. 2015). We used a developmental threshold of 10 degrees as in Cayton et al. (2015), and calculated GDD over the period from January 1 through July 1 of each year to represent the time before most butterflies eclosed (for details, see Appendix S1). For each population metric (abundance, mean day of activity, flight period), we fit a simple linear regression with GDD as the predictor. We also calculated 95% confidence intervals for the slope using parametric bootstrapping, and the proportion of p values that were less than 0.05 among these bootstrapped model fits.

### Results

#### Estimation of phenology metrics

For this univoltine butterfly population, the Gaussian curve provides a visually satisfying fit, with the model reasonably fitting years with many surveys (Fig. 1 A-F) and those with few (2018-2020, Fig. 1G-I). We found no overall indication of asymmetry in activity when fitting our residuals with a linear model (slope = 0.085, p=0.57), and our fitted cubic spline showed no notable deviations from a linear model (estimated degrees of freedom for the smoothing term was 1, suggesting a straight line is the best fit). Estimates of our three metrics (Abundance index, flight period, and day of peak activity) were generally very precise, with notable exceptions for 2019 and 2020, years with only a few non-zero survey counts (Figs. 2A-C). We also see a strong correspondence between our abundance index and population estimates from the capture-recapture study (R^2^ = 0.94) (Fig. 3A). These data also demonstrate the bias of first and last dates of observations in relation to population size (Fig. 3C-D); compared to 0.1 and 0.9 quantiles estimated from annual Gaussian curves, years with smaller populations had later first observations and earlier last observations.

**Figure 2:**
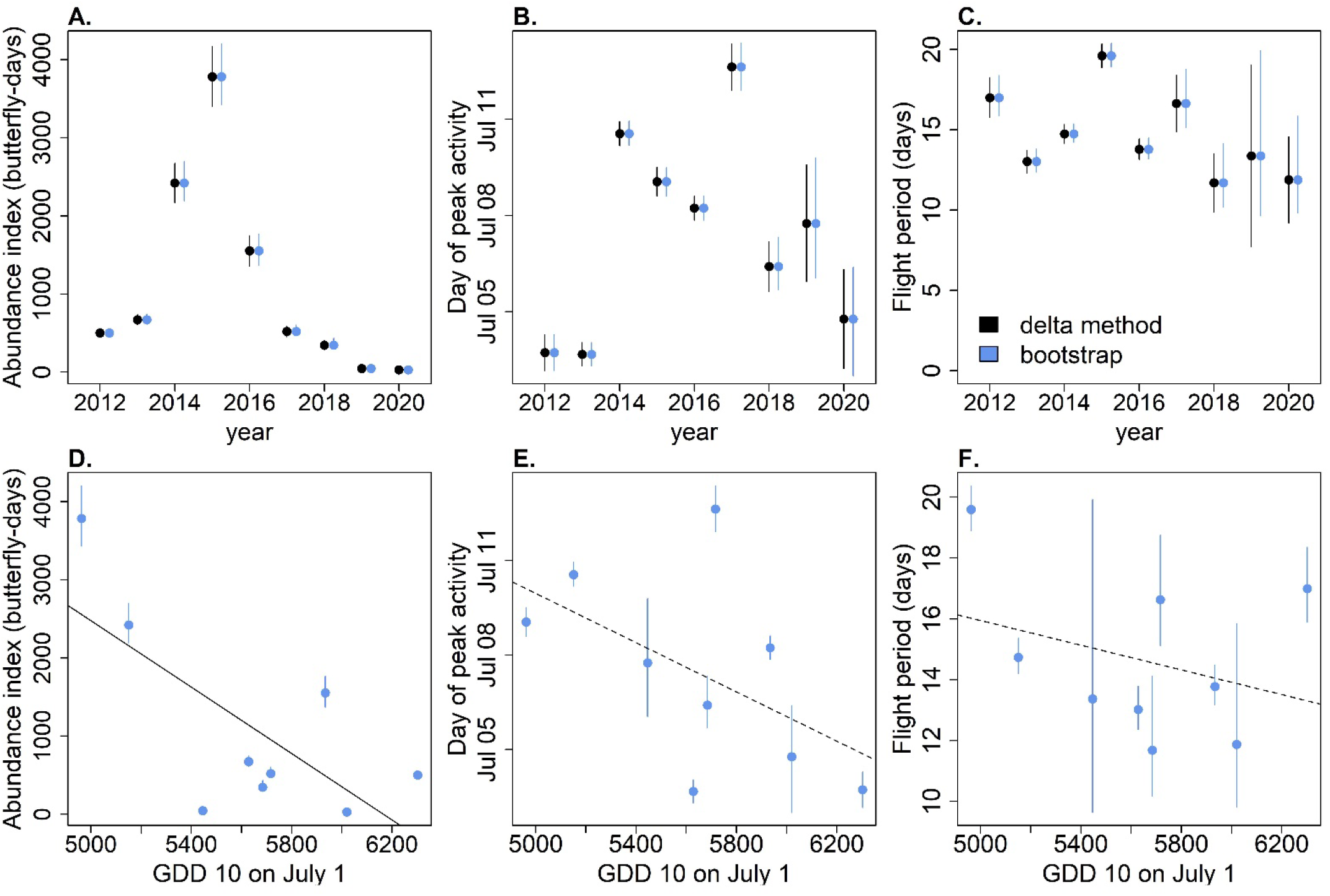
Fitted population metrics: (A-C) abundance index (A), mean day of activity (B), flight period (C) across years (sequentially). Bars show +/− 1 standard error, calculated using the delta method (black) or parametric bootstrapping (blue). (D-F) Comparison of same population metrics with the Growing Degree Days (GDD) on July 10 for each year. Solid and dashed black lines show best-fitting model for significant (solid) and non-significant (dashed) linear regression. In all panels, parametric bootstrap standard errors are based on 0.16 and 0.84 quantiles of bootstrap samples (quantiles corresponding to +/− 1 standard error).

**Figure 3:**
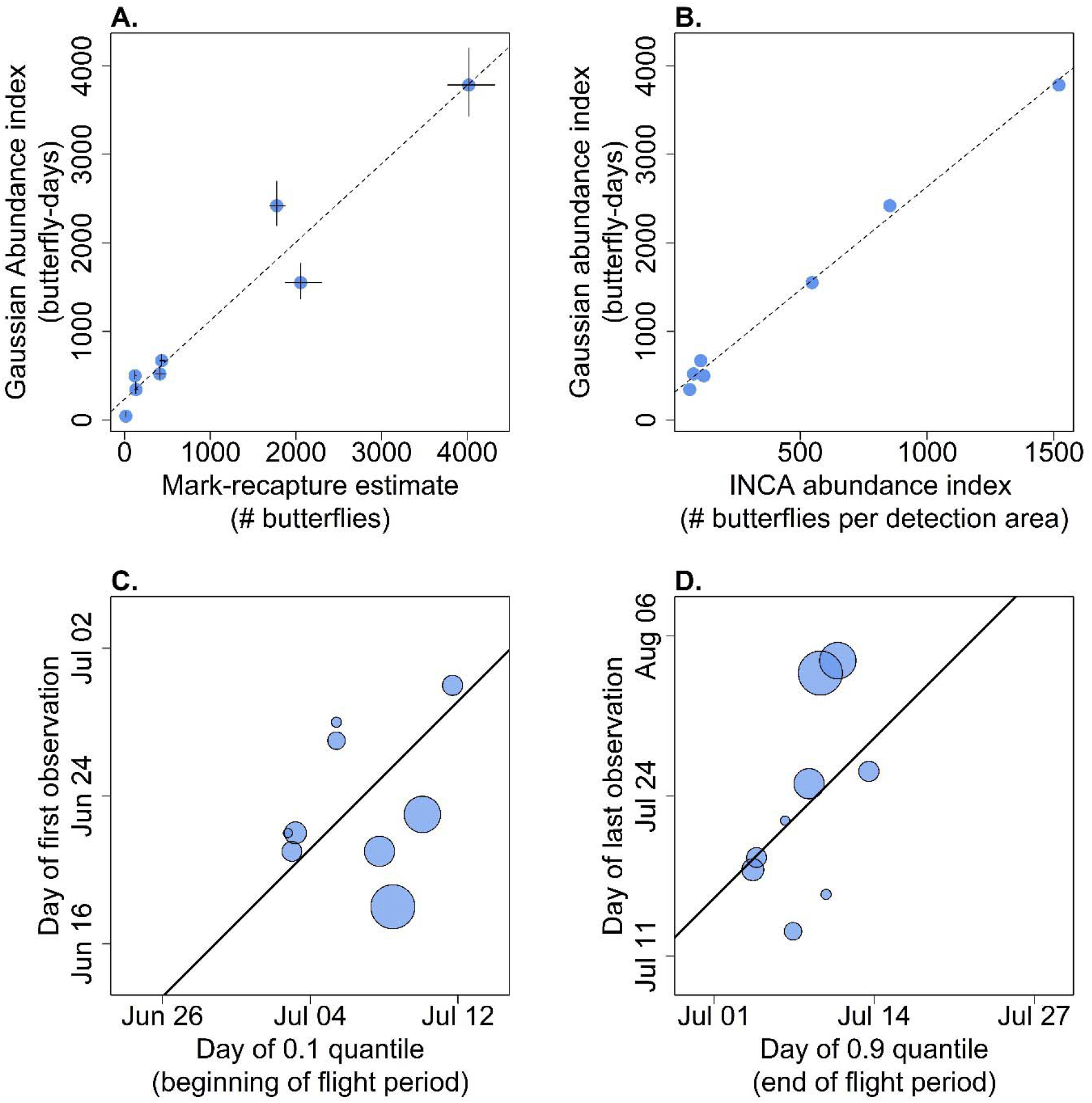
Comparing derived estimates from GLM approach to other commonly used approaches (A) Comparison of GLM estimated abundance index to mark-recapture estimates. Black lines show +/− 1 standard error of each metric, dashed line shows best-fitting curve (R^2 = 0.94). (B) Comparing CLM estimated abundance index with INCA estimated abundance index (INCA is an implementation of the Zonneveld model) (R^2 = 0.99). (C-D) Comparison of quantile-based estimates of onset (C) and end (D) of activity to day of first (C) and last (D) observation for each year, with point size scaled by GLM estimated population index (square-root scale). Black lines show a 1:1 line going through the mean of observed values. As expected, years with smaller population sizes show less extreme observations (i.e., points above (C) and below (D) the 1:1 line).

#### Comparison to INCA fits

Without independent estimates of mortality, INCA fit only 3 of the 9 years of data (Table S1). Using published mortality estimates (Brown and Crone 2016) as an informative prior probability distribution, INCA was able to fit more years of data (7 of the 9), but still failed to fit 2019 and 2020. For the years in which the informed INCA model fit, there was a very strong correspondence between the informed INCA fit and the Gaussian fit, with an R-squared of 0.996 (population abundance indices) and 0.883 (day of peak activity) (Fig. 3B).

#### Evaluating an environmental driver

Temperature had a strong association with population abundance index, with warmer years associated with smaller population indices (estimate slope: −2.12; bootstrapped 95% CI of slope of N vs GDD: [−2.3087, −2.0598]; across our bootstraps, this was almost always significant (p<0.05 in 99.6% of bootstraps) (Fig. 3D)). Mean day of activity was consistently earlier in warmer years (estimated slope: −0.004; bootstrapped 95% CI of slope of *μ* vs. GDD: [−0.0065, −0.0024]), but it was rarely statistically differences in flight period (estimated slope: −0.002; bootstrapped 95% CI of *σ* vs. GDD: [-significant (p<0.05 only 6 out of 10,000 times) (Fig. 3E). Temperature was not associated with 0.0023, 0.0007], p<0.05 0 of 10,000 times) (Fig 3F).

## Discussion

In this paper, we show how a Gaussian curve can be fit to insect count data using familiar methods for linear models, and that it allows us to estimate abundance and phenology even for years of sparse data where other methods can fail. We hope this approach provides a much-needed tool for ecologists trying to study insect decline or the phenology and dynamics for at-risk species (or species that have sparse count data for other reasons). There is a particular need for tools like this given the growing interest in documenting and understanding insect decline; our ability to do so is in large part limited by available data and methods (Didham et al. 2020). We are not the first to use Gaussian curves to fit count data (e.g. Lindén and Mäntyniemi 2011, Dennis et al. 2015, Oke et al. 2019, Stewart et al. 2020), but past implementations have required custom coding and more advanced knowledge of statistical computing.

The combination of a simple mathematical form (3 parameters) and the robust fitting algorithms associated with linear models allows Gaussian models to estimate phenology and abundance even in years with relatively few observations (see, e.g., Fig 1H). Recent studies of butterfly (Hodgson et al. 2011, Wepprich et al. 2019, Stewart et al. 2020) and bee (Stemkovski et al. 2020) populations have generally been restricted to relatively abundant species by the needs of their more data-hungry methods. These more flexible analytical tools like GAMs provide more detailed information about activity curves, but at the cost of requiring sufficient data to differentiate between the many possible shapes those more flexible curves can take. In contrast, while the Guassian curve is constrained in shape and cannot capture complex activity curves, we are consistently able to fit curves with only 3 non-zero surveys (XXX unpubl. simulations). Our goal is not to replace existing tools, which often provide more detailed information than a Guassian curve can, like capturing multimodality (e.g. GAMs) or measuring asymmetry and linking it to biological processes (e.g. the Zonneveld model). Rather, we want to “unlock” data sets which were previously unusable either because the observations were too sparse for other methods, or interested parties did not have the computational statistics background needed to fit more complex models. We explain and demonstrate this method assuming a simple data structure (e.g. multiple years, but one species and one site). With hierarchical data (e.g. multiple sites, multiple species), this method can be expanded upon to fit separate curves for statistical unit (e.g. each year of each site) (Bonoan et al., *in review).*

Comparing fits and estimates of the Baltimore checkerspot butterfly using our Gaussian method, the INCA implementation of the Zonneveld model, and capture-recapture tools demonstrate the value of our approach. Without outside information, the INCA model fit only one third of our 9 years of data, and even with the inclusion of an independent estimate of mortality rates, INCA failed to fit the two years with the lowest estimated abundance (Table S1). However, for years when we could fit the data using the Zonneveld/INCA model informed by independent estimates of mortality, we see a very strong correspondence between INCA and Gaussian estimates of population abundance indices (R^2 = 0.99) (Fig. 3B), suggesting that our proposed method is a useful and comparable alternative to INCA when data are sparse. We also see a tight correlation between the abundance index of the Gaussian model, and capture-recapture estimates of population size calculated separately from the same data (Fig. 3A), which suggests that abundance estimates are unbiased. This correlation compares favorably with other methods of fitting transect data; Haddad et al. (2008) found no correlation between mark-recapture estimates of population size and population size estimated using the Zonneveld model. However, the fact that we find a 1:1 match of *N* (an index that reflects longevity as well as abundance) and capture-recapture estimates is likely coincidental. By chance, our capture probability during surveys was ≈ 0.15 (XXX and YYY unpubl.), and the apparent survival of Baltimore checkerspot butterflies at our site is 0.844/day (Brown and Crone 2016); these values mean that in our example, the capture probability - by chance - exactly cancelled out the fact that *N* is actually in units of “butterfly days”.

While it is becoming increasingly rare, objectively problematic metrics for phenological patterns such as first or last observations are still used by at least some ecologists (e.g. Fric et al. 2020, Colom et al. 2020). For many types of data sets, observations of first and last events are known to be biased, as the day of first or last observation depends in part on population size and detectability (Van Strien et al. 2008, Miller-Rushing et al. 2008, Inouye et al. 2019). Of course, sometimes data limitations constrain analysis to only use first or last metrics, especially when comparing with historic data sets (e.g. Heberling et al. 2019). However, in many cases ecologists have much more complete data, and should not be limited to using problematic phenological metrics. This point has been made thoroughly in other studies; as expected, for the Baltimore Checkerspot we see consistent biases in first and last date observed based on population size (Fig. 3C-D). As an alternative to problematic metrics, fitting Gaussian curves may be a reasonable first step for many ecologists interested in describing phenology. Early and late quantiles (e.g. 0.1 and 0.9, as in Jonzén et al. (2006) and Michielini et al. (2020), or 0.05 and 0.95 as in Stemkovski et al. 2020) can easily be calculated from estimated μ and σ, and are unbiased analogs to represent the early and late parts of the activity season (cf. Bonoan et al., *in review*).

We demonstrated how our approach can be used to link population-level patterns with environmental (or other) drivers. In doing so, we found a significant negative relationship between growing degree day (GDD) and abundance indices, and a non-significant pattern of earlier activity in warmer years that was consistent across bootstraps. These results are largely consistent with the patterns found in other studies. Warmer temperatures have led to earlier activity for butterfly species in the UK (MacGregor et al. 2019), Spain (Stefanescu et al. 2003, Stewart et al. 2020), and Ohio (Cayton et al. 2015), and studies have found that in recent decades butterflies have advanced their phenology in the UK (MacGregor et al. 2019) and across the northern hemisphere (Parmesan 2007). The relationship between temperature and abundance across studies is more complicated. Studies have found warmer temperatures leading to higher population abundance in most butterfly species in the UK (Roy et al. 2001) and a mixture of butterfly abundance responses to temperature in Spain (Stewart et al. 2020). In contrast, Isaac et al. (2011) found butterfly density in England was generally lower in regions with higher temperatures, and Colom et al. (2020) found warmer summers were associated with smaller butterfly populations on the Spanish island of Menorca. In Massachusetts USA, butterfly populations near their species’ northern range limits are generally increasing, and populations near their species southern range limits are generally decreasing (Breed et al. 2012, Michielini et al. 2020).

Gaussian curves are only well-suited to represent data that is unimodal and approximately symmetric. For many phenological events, the assumption of symmetry may be a reasonable approximation (see Fig 1, and Stewart et al. 2020), although this is of course a hypothesis that could be explored depending on the goals of an analysis (in our analysis of Baltimore checkerspot, residuals did not indicate skew). Multimodal distributions may be more problematic. For multivoltine insects, Generalized Additive Models (GAMs) (e.g. Knudsen et al. 2007, Moussus et al. 2009, Hodgson et al. 2011, Newson et al. 2016, Stemkovski et al. 2020) have been used to capture changes in phenology over time. Although they are not described by a parametric equation, features like the onset (0.1 quantile) or end (0.9 quantile) can be extracted from GAMs numerically (cf. Stemkovski et al. 2020). Another approach to evaluating phenological events without assuming a particular distribution is quantile regression (Cade and Noon 2003, Koenker 2019), which has been used in several studies of bird migration (e.g. Gordo et al. 2013, Barton and Sandercock 2018), and occasionally for Lepidoptera (Gimesi et al. 2012, Michielini et al. 2020). Like GAMs and GLMs, quantile regression shares the property of drawing on well-established and well-validated statistical approaches, rather than developing new ones.

Understanding trends in abundance has long been a goal of both population ecology and conservation management, and this has become all the more urgent with observed and suspected population declines in a wide range of species, particularly insects. Similarly, because phenological shifts are one of the most conspicuous signs of climate change, there is growing interest in their causes and consequences. We expect that the widespread interest in abundance and phenology will continue to lead to a growing number of new methods for interpreting patterns in count data. At the same time, not every new method is guaranteed to work for all (or even most) data, and custom-coding for every question can be error-prone, time consuming, and intimidating to many ecologists. We encourage ecologists to be aware of well-established existing methods, and provide the linearized Gaussian model as a simple tool for unlocking previously-inaccessible sparse data sets.

## Declarations

This work was conducted with support from NSF DEB 19-20834 and the DOD SERDP program (RC-2700). We thank many XXXX lab members who participated in Baltimore checkerspot data collection, especially YYY YYYY and ZZZZ ZZZZZZZZ, and Brian Inouye, Jessica Forrest, and two anonymous reviewers for helpful suggestions on the manuscript. Data and code will be available on Dryad.

---

**Table S1:** Comparing ability of GLM and INCA models to fit the Baltimore checkerspot data. “Non-zero counts” represents the number of survey days in that year with at least one butterfly observed; “Gaussian fits?” shows whether or not our GLM approach fit the data”; “INCA fits?” shows whether or not the Zonneveld method implemented in the INCA program fit the data (using default settings); “INCA+ fits?” shows the same results, but when INCA is given mortality information independently determined from Brown and Crone 2016. We see that INCA struggles to fit much of our data, INCA+ fits most of our data, and the GLM approach fits all of our data.

| Year | Non-zero counts | INCA fit? | INCA+ fit? | Gaussian fit? |
| --- | --- | --- | --- | --- |
| 2012 | 23 | **<u>No</u>** | Yes | Yes |
| 2013 | 20 | Yes | Yes | Yes |
| 2014 | 20 | **<u>No</u>** | Yes | Yes |
| 2015 | 18 | **<u>No</u>** | Yes | Yes |
| 2016 | 15 | **<u>No</u>** | Yes | Yes |
| 2017 | 11 | Yes | Yes | Yes |
| 2018 | 6 | Yes | Yes | Yes |
| 2019 | 5 | **<u>No</u>** | **<u>No</u>** | Yes |
| 2020 | 4 | **<u>No</u>** | **<u>No</u>** | Yes |

## Footnotes

1 In ecology. The Gaussian mixture model (Proia et al. 2016) has been used several times in publications on highway maintenance and once on diagnostics of ventricular septal defects.

